# The structure of monkeypox virus 2’-O-ribose methyltransferase VP39 in complex with sinefungin provides the foundation for inhibitor design

**DOI:** 10.1101/2022.09.27.509668

**Authors:** Jan Silhan, Martin Klima, Dominika Chalupska, Jan Kozic, Evzen Boura

## Abstract

Monkeypox is an emerging, rapidly spreading disease with pandemic potential. It is caused by the monkeypox virus (MPXV), a dsDNA virus from the *Poxviridae* family, that replicates in the cytoplasm and must encode for its own RNA processing machinery including the capping machinery. Here, we present the crystal structure of its 2’-O-RNA methyltransferase (MTase) VP39 in complex with the pan-MTase inhibitor sinefungin. A comparison of this 2’-O RNA MTase with enzymes from unrelated ssRNA viruses (SARS-CoV-2 and Zika) reveals a surprisingly conserved sinefungin binding mode implicating that a single inhibitor could be used against unrelated viral families.

## Introduction

Monkeypox is a disease caused by the monkeypox virus (MPXV), an Orthopox virus belonging to the *Poxviridae* family, clinically resembling the eradicated smallpox^1^. Until recently, its occurrence was limited to central and western Africa where its natural reservoir consists of rodents and primates^2^; however, it can also infect humans, and the mortality rate is estimated to be 3 - 6%. This rate is lower than the deadly smallpox, but much higher than COVID-19 that is caused by SARS-CoV-2. Monkeypox has become increasingly spread across the globe with number of cases exponentially growing reaching somewhat alarming 54900 cases, and it has been reported in some 100 countries^3^. It is no wonder that the prospective of another viral pandemic alarmed the public health authorities as well as the general public.

MPXV is a dsDNA virus; however, it replicates in cytoplasm, implying that it must encode for DNA and RNA replication machinery^4^ because the human enzymes are located in the nucleus. Besides the DNA-dependent DNA polymerase and the RNA-dependent RNA polymerase, it also encodes for the RNA capping machinery. In an early stage of cellular infection, the RNA cap is important for the initiation of translation of the viral RNA (vRNA)^5,6^. This cap plays an important role in the innate immunity as uncapped RNA is recognized by the innate immunity IFIT and RIG-I sensors ^7,8^. The cap is also important for stability of the mRNA, thus the poxviruses encode for decapping enzymes to prevent accumulation of dsRNA later during infection which would induce the innate antiviral response^9^. Indeed, protection against innate immunity is of the uttermost importance for the MPXV, and it also encodes poxin, an enzyme that blocks the cGAS-STING pathway triggered by the presence of dsDNA in cytoplasm^10^.

Chemically, the mRNA cap is a structure where an N7-methylated guanosine is linked via a triphosphate to the 5’ end of the RNA yielding m7GpppRNA, which is also referred to as cap-0. In the fully matured cap (cap-1), the first nucleotide is also methylated at the 2’-O position of the ribose moiety. In the *Poxviridae* family, cap-0 is synthesized in cytoplasm by the heterodimeric capping enzyme that has RNA 5’ triphosphatase (RTPase), guanylyltransferase (GTase) and (guanine-N7)-methyltransferase (MTase) activity^11^. The addition of another methyl group to the 2’-O position of the adjacent ribose converts cap-0 to cap-1. This step is also important in preventing an innate immunity response^12^ and is catalyzed by the MPXV 2’-O MTase VP39^13^. Here, we present the structure of the MPXV VP39 in complex with the pan-MTase inhibitor sinefungin. The structure reveals the mechanism of VP39 inhibition and comparison of this structure to the 2’-O MTases from unrelated viruses SARS-CoV-2 and Zika has important implications in the design of pan-antivirals based on MTase inhibitors.

## Results

### Overall structure and enzymatic characterization of the MPXV VP39 MTase

To obtain the structure, we expressed and purified recombinant MPXV VP39 in *E. coli*. and performed screening of the crystallization conditions. Initial crystals were of poor quality but diffraction quality crystals were obtained upon a few rounds of seeding. The crystals diffracted to 2Å resolution and belonged to the orthorhombic P2_1_2_1_2_1_ space group. The structure was solved by molecular replacement and refined to R_work_ = 20.81% and R_free_ = 23.24% and good geometry (detailed in Table 1). The structure revealed a mixed α/β fold that resembles the Rossmann fold. A central β-sheet is composed of β-sheets β2-β10 that adopt the shape of the letter *J*. Interestingly, this arrangement was also observed for the SARS-CoV-2 2’-O MTase nsp16^14^. The central sheet is stabilized from one side by helices α1, α2, α6 and α7, and by α3 and α7 from the other. These opposite sides of the *J*-like β-sheet are connected by α 5, β1 and β11 (Figure 1A, B).

**Table 1.**
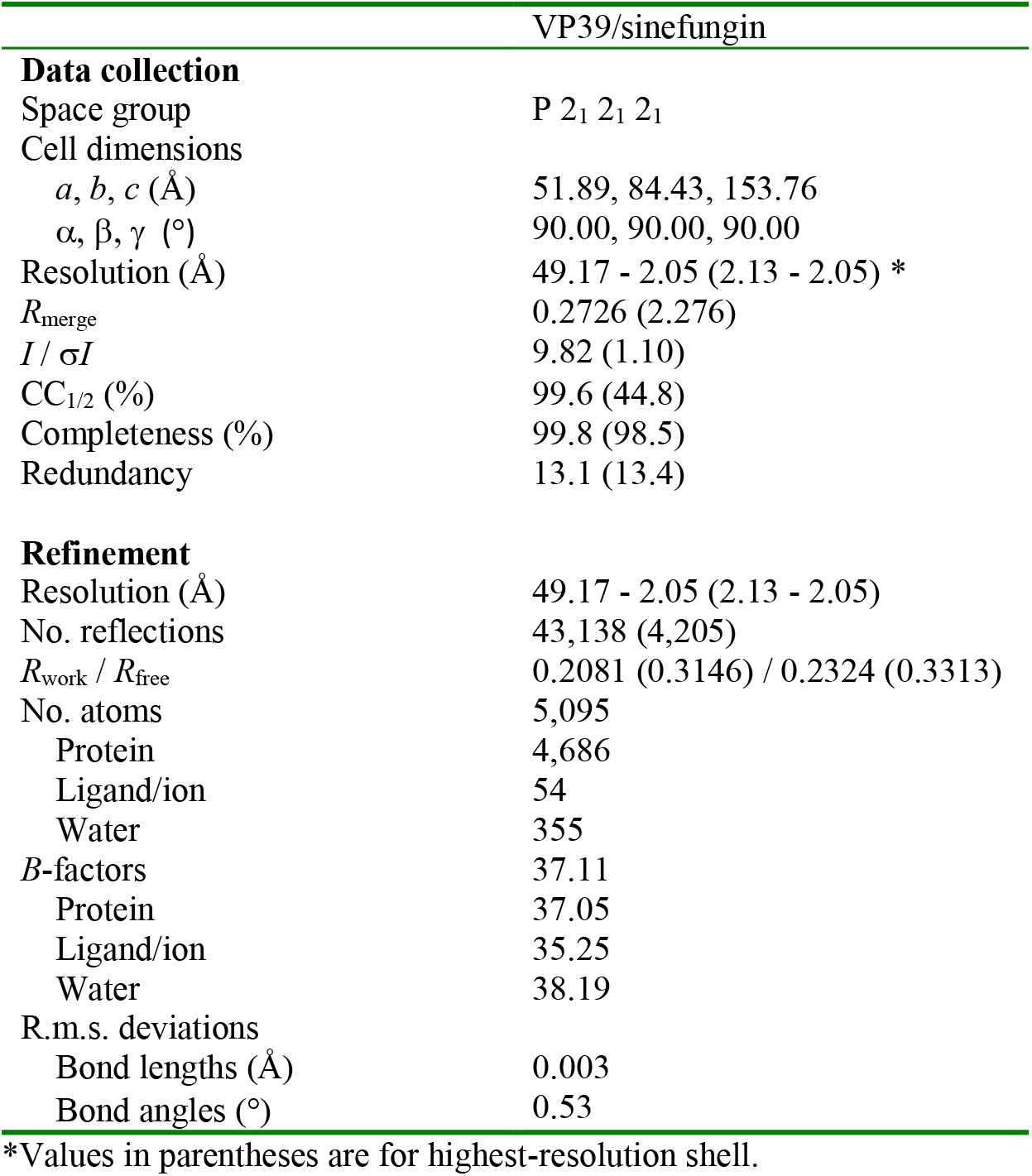
Data collection and refinement statistics.

**Figure 1:**
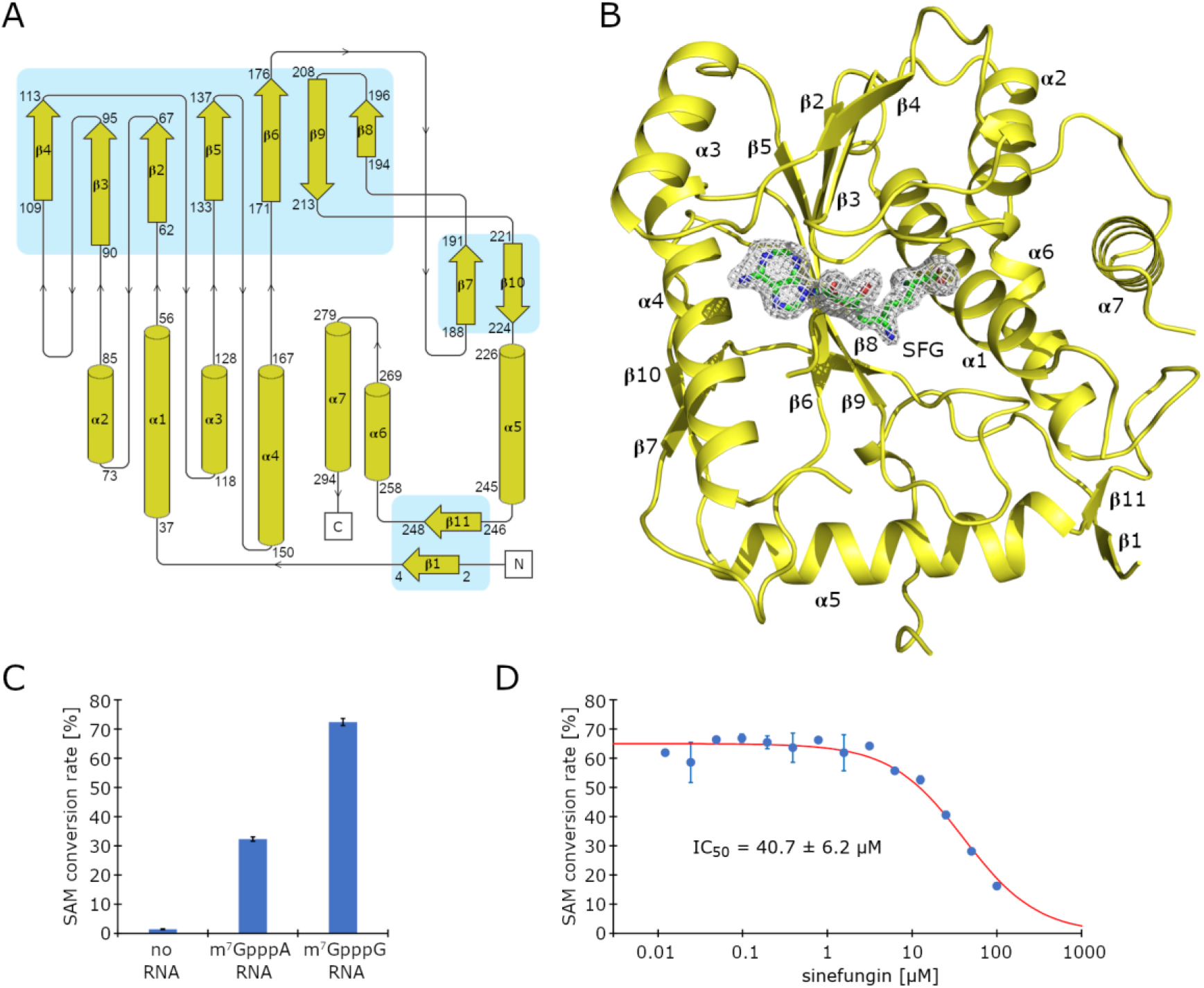
Crystal structure of the monkeypox virus VP39 methyltransferase in complex with sinefungin. A) Topology plot of the monkeypox virus VP39 protein. B) Overall fold of the VP39 protein in complex with sinefungin. The protein backbone is shown in cartoon representation and depicted in yellow, while sinefungin is shown in the stick representation and colored according to its elements: carbon, green; nitrogen, blue; oxygen, red. The unbiased Fo-Fc omit map contoured at 3σ is shown around sinefungin. C) Methyltransferase activity of the VP39 protein with capped RNA substrates used as indicated. Data points are presented as mean values ± standard deviations (n = 3). D) Inhibition of the MTase activity of the VP39 by sinefungin. Data points are presented as mean values ± standard deviations from triplicates. IC_50_ value is presented as the best fit ± 95%confidence interval of the fit.

We tested two substrates that differed in the penultimate base (m7GpppA-RNA vs. m7GpppG-RNA) to verify the enzymatic activity of our recombinant VP39 enzyme. Both substrates were accepted; however, the guanine base was clearly preferred in the penultimate position (Figure 1C). That is in contrast to coronaviruses where adenine is preferred at this position^15^. We also verified that sinefungin was able to inhibit the VP39 MTase; under the conditions used (detailed in M&M), it exhibited IC_50_ = 40.7 ± 6.2 μM (Figure 1D).

### Analysis of the VP39-sinefungin interaction

The electron density for sinefungin was immediately visible upon molecular replacement in the central pocket of the VP39 MTase (Figure 1). Sinefungin occupied the SAM binding pocket (Figure 2A). Its adenine moiety lies within a deep canyon lined with hydrophobic sidechains from residues Phe115, Val116, Val139 and Leu159 and makes two hydrogen bonds with the backbone of Val116 (Figure 2B). The ribose and amino acid moieties are also involved in hydrogen bonding: the ribose ring directly binds the Asp95 and Arg97 via its hydroxyl groups while the amino acid moiety forms hydrogen bonds with the sidechains of Gln39 and Asp138 (Figure 2B).

**Figure 2:**
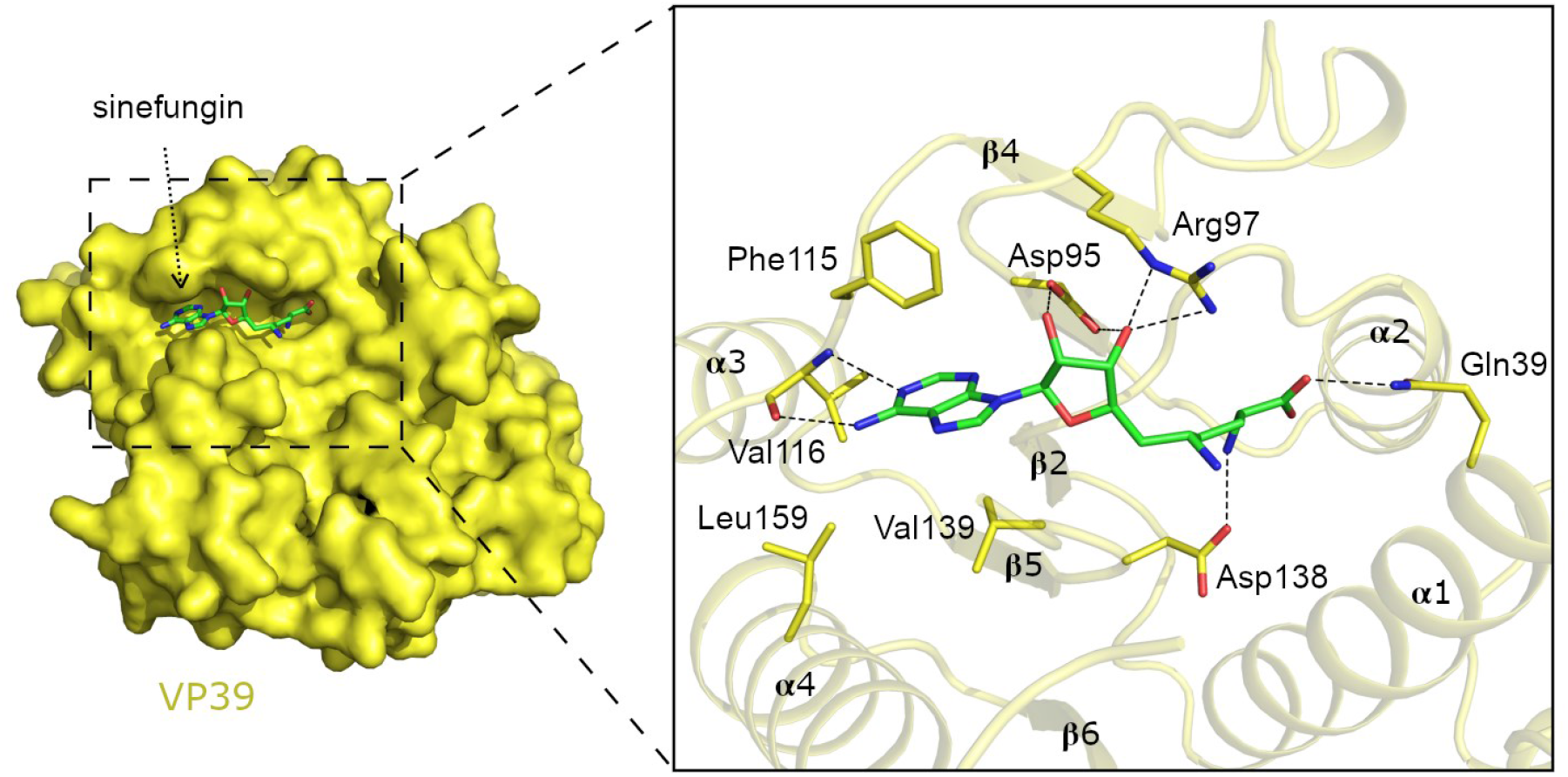
Sinefungin recognition by the monkeypox virus VP39 methyltransferase. Detailed view of the sinefungin binding site. Sinefungin and side chains of selected VP39 amino acid residues are shown in the stick representation with carbon atoms colored according to the protein/ligand assignment and other elements colored as in Fig 1B. Selected hydrogen bonds involved in the VP39-sinefungin interaction are presented as dashed black lines.

Based on the previously solved crystal structure of vaccinia VP39 in complex with RNA^16^, we constructucted a model of a ternary complex sinefungin:RNA:VP39 to illustrate the molecular mechanism of sinefungin’s action (Figure 3). It efficiently protects the 2’-O-ribose position; its amino group is in the close vicinity of the 2’ ribose hydroxyl group where the sulphur atom of SAM would be located otherwise.

**Figure 3:**
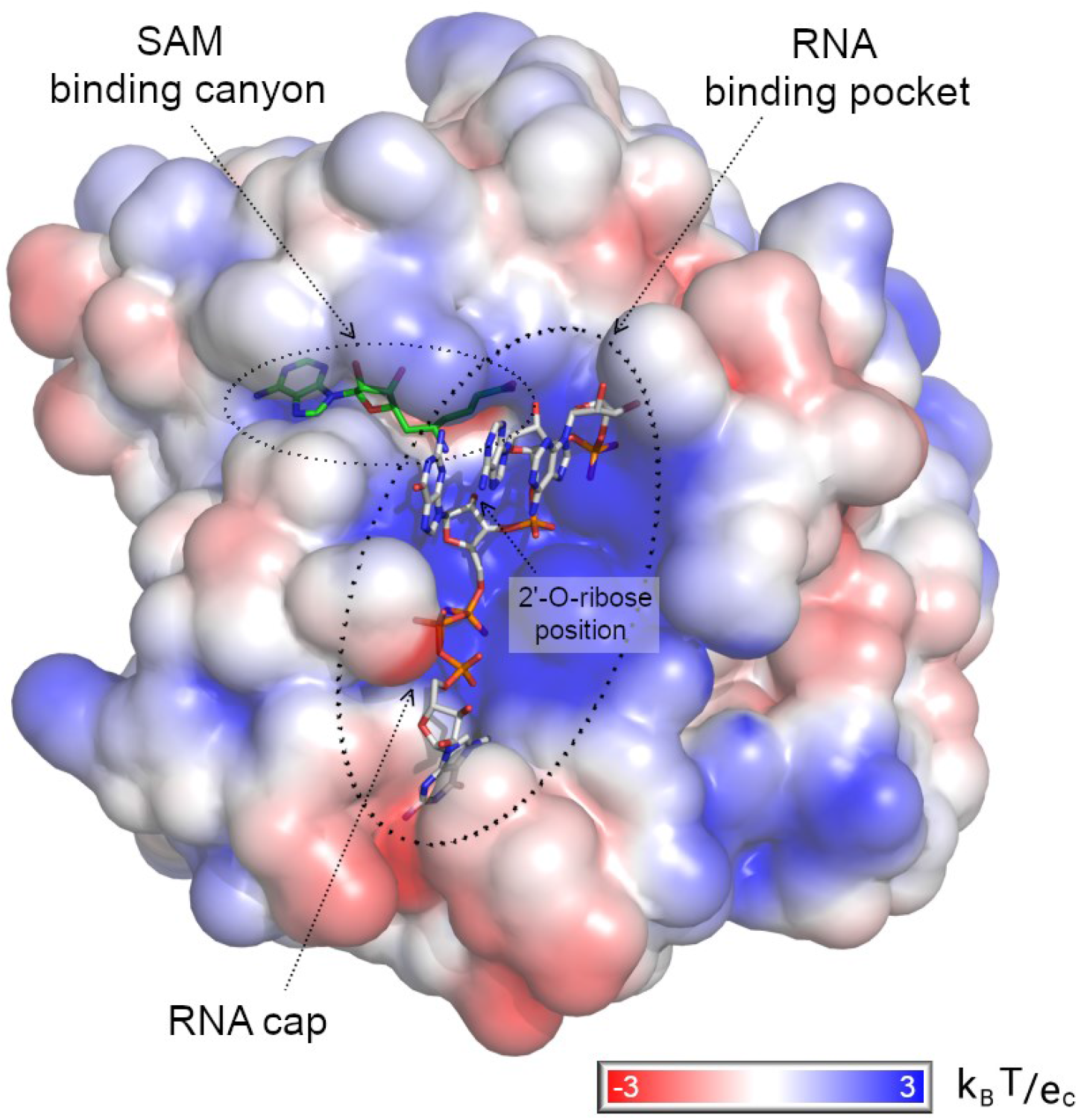
Model of RNA recognition by the monkeypox virus VP39 methyltransferase. Surface of the VP39 protein is colored according to the electrostatic surface potential. The m7GpppG-capped RNA was modeled by structural alignment using the crystal structure of the vaccinia virus VP39 protein in complex with SAH and m7GpppG-capped RNA (pdb entry 1av6) as a template. Sinefungin and m7GpppG-capped RNA are shown in the stick representation and colored according to the elements: carbon, green (sinefungin) or white (m7GpppG-capped RNA); nitrogen, blue; oxygen, red; phosphorus, orange.

### Implications for inhibitor design

The SAM binding canyon has two ends: one borders with the RNA binding pocket and serves to position SAM for the methyltransferase reaction; the other end adjacent to the adenine base of sinefungin is apparently unoccupied (Figure 3). However, a close inspection of this location reveals a network of water molecules that are interconnected by hydrogen bonds and also bond the adenine moiety and residues Val116, Glu118 and Asn156 (Figure 4). Compounds based on the sinefungin scaffold bearing a moiety that would displace these waters and interact with Val116, Glu118 and Asn156 directly could be an exceptionally good binder because displacement of water molecules can have a favorable entropic effect^17,18^.

**Figure 4.**
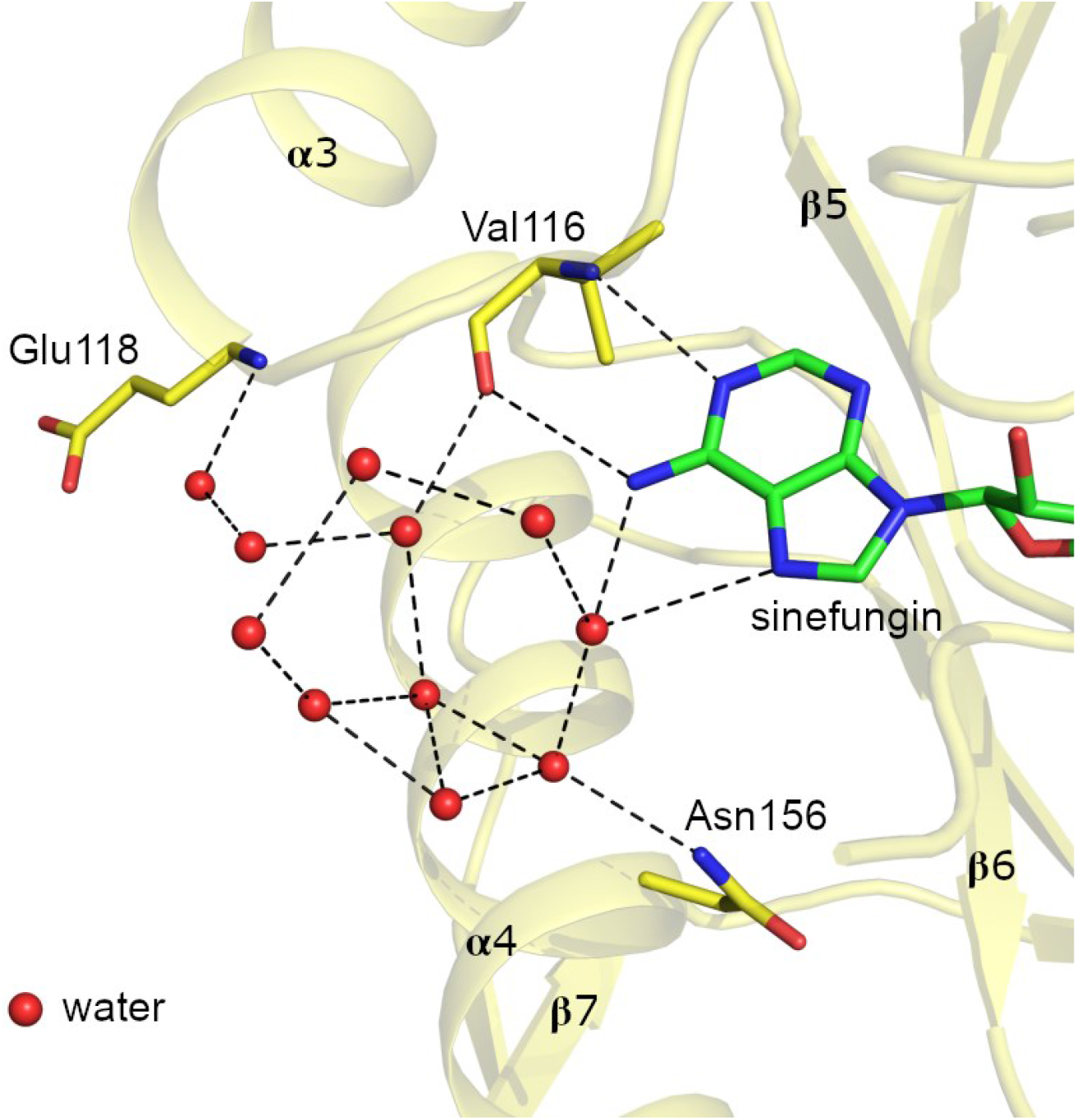
A network of coordinated water molecules in the vicinity of the sinefungin binding groove. Detailed view of the water molecules coordinated next to the adenine base of sinefungin. Sinefungin and the side chains of selected VP39 amino acid residues are shown in the stick representation with carbon atoms colored according to the protein/ligand assignment and other elements colored as in Fig 1B. Water molecules are shown as red spheres; selected hydrogen bonds are presented as dashed black lines.

Next, we compared the catalytic sites of VP39 with the catalytic sites of 2’-O-ribose MTases from unrelated medically important viruses, namely SARS-CoV-2 and Zika. The resemblance of the SAM binding sites is remarkable in both cases. Sinefungin is virtually in the same conformation in MPXV VP39, SARS-CoV-2 nsp16 and Zika NS5 (Figure 5). The catalytic tetrad (Lys41, Asp138, Lys175, Glu218 for MPXV) is absolutely conserved among these unrelated viruses including the conformation of these residues. However, differences do exist in the binding mode of the nucleobase as well as the sugar ring. The adenine base is bound to the backbone of Val116 (Figure 2) and held in place by a complex water network in the case of VP39 (Figure 4), while in SARS-CoV-2 it is held by the side chain of Asp114 (Figure 5A) and in Zika by the side chain of Asp131 (Figure 5B). The binding mode of the ribose ring is virtually identical between VP39 and SARS-CoV-2 nsp16, as both of these viruses utilize an aspartate residue (Figure 5A). However, in Zika it is held in place by His110 and Glu111. Interestingly, all these viruses use an aspartate residue to interact with the amino group of sinefungin’s amino acid moiety, further highlighting the similarity of the SAM binding site between a dsDNA virus and two distinct +RNA viruses.

**Figure 5:**
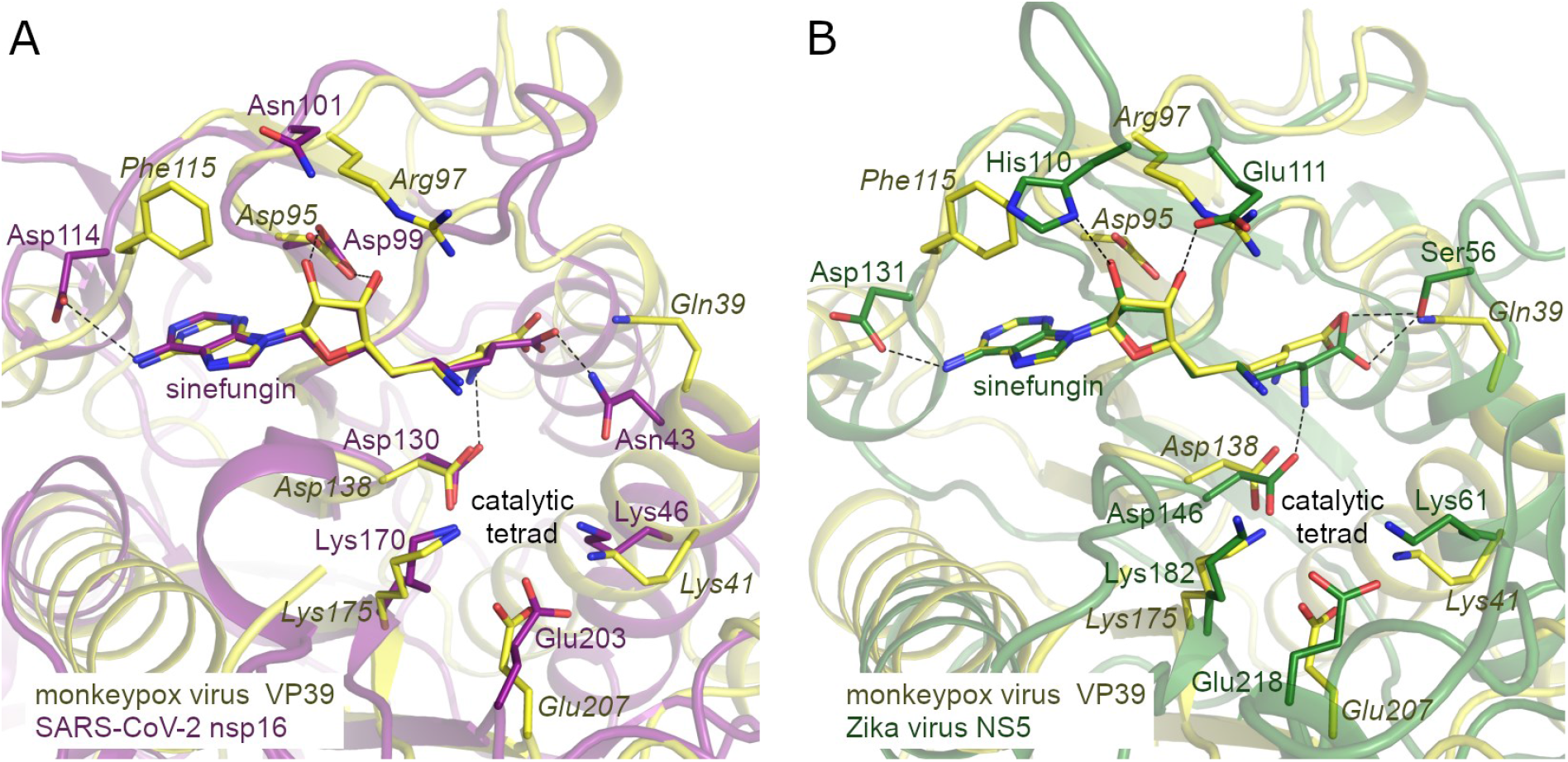
Structural alignment of the methyltransferase active sites of monkeypox virus VP39 with SARS-CoV-2 nsp16 and Zika virus NS5. Crystal structures of SARS-CoV-2 nsp10/nsp16 (A, pdb entry 6yz1) and Zika virus NS5 methyltransferase domain (B, pdb entry 5mrk) were used for the alignment. Protein backbones are shown in the cartoon representation and depicted in yellow (monkeypox virus VP39), magenta (SARS-CoV-2 nsp16), and green (Zika virus NS5). Sinefungin and the side chains of selected residues are shown in the stick representation with carbon atoms colored according to the protein assignment and other elements colored as in Fig 1B.

## Discussion

Viruses are a global threat to human health as illustrated by the recent COVID-19 pandemic. Even viral diseases with mortality rates ~ 1% can kill millions of people if they spread globally. Monkeypox is already present on every continent, its spread was very likely facilitated by cessation of the vaccination against smallpox in the 1980s since the smallpox vaccine is effective against the monkeypox (~80% efficiency)^19^. With an estimated mortality rate of 3 - 6%, the MPXV has the potential to cause a devastating pandemic^20^. On the other hand, the effective FDA-approved vaccine JYNNEOS^21^ is available, as is the FDA-approved drug - tecovirimat^22^. Encouragingly, MPXV is a dsDNA virus, and the mutation rate of DNA viruses is much lower than that of RNA viruses, such as SARS-CoV-2 or Zika. Nonetheless, it would be wise to develop more drugs to ensure their availability and efficiency.

RNA MTases are present in multiple unrelated viral families, including both RNA and DNA viruses^23^; yet, none of the FDA approved drugs target a viral MTase. Most current antivirals are inhibitors of polymerases, integrases or proteases. In fact, only a handful of inhibitors of viral MTases, mostly targeting flaviviruses^24,25^, were known prior to the COVID-19 pandemic. However, recently many inhibitors of SARS-CoV-2 MTases were reported by us and others^26–30^.

It remains to be established if inhibitors of MPXV MTases are viable antiviral targets; however, the high conservation of VP39 across orthopoxviruses suggests an important role in the viral lifecycle. In addition, the similarity of the dsDNA virus VP39 SAM binding site with the 2’-O MTases from two +RNA viral families (flaviviruses and coronaviruses, Figure 5) suggests that the development of a pan-antiviral MTase inhibitor should be feasible, especially since the catalytic tetrad is absolutely conserved. However, to obtain specific antiviral inhibitors that will not interfere with the human RNA 2’-O MTases, it might be necessary to take advantage of specific features of the viral enzymes, such as a large hydrophobic pocket above the adenine base that is present in coronavirus nsp16^28,31^ or the complex water network adjacent to the adenine base in the case of MPXV VP39. Nonetheless, inhibitors of viral MTases are urgently needed to further elucidate the role of these enzymes during viral infection.

## Materials and Methods

### Protein expression and purification

The gene for the monkeypox (USA-May22 strain) viral MTase VP39 was codon-optimized for expression in *Escherichia coli* and commercially synthesized (Azenta). Subsequently it was cloned into a pSUMO (with 8xHis-SUMO tag) using restriction cloning (BamHI and NotI sites). The recombinant protein was purified according to our established protocols for viral methyltransferases^32^. Briefly, *E. coli* BL21(DE3) cells were transformed with the VP39 expression plasmid and incubated in LB medium supplemented with ZnSO4 (10 μM) and ampicillin (100 μg/ml) and cultured at 37°C. When the cells reached optical density [OD] = 0.6 the production of VP39 was induced by the addition of IPTG (isopropyl-b-D-thiogalac-topyranoside) (250 μM) and the temperature was lowered to 20°C for 15 h. The cells were pelleted by centrifugation and resuspended in lysis buffer (25 mM Tris, pH 8, 300 mM NaCl, 30 mM imidazole pH = 8.0, 3 mM b-mercaptoethanol) and lysed by sonication. The clarified lysate was subjected to affinity chromatography on Ni^2+^ agarose resin (Macherey-Nagel) and washed extensively with lysis buffer. The recombinant protein was eluted with lysis buffer supplemented with 300 mM imidazole pH = 8.0. The 8xHis-SUMO tag was cleaved by Ulp1 protease during dialysis against the lysis buffer. Subsequently the Ulp1 and His8x-SUMO tag were removed by a second round of affinity chromatography. Unbound fraction was concentrated and loaded on HiLoad 16/600 Superdex 75 gel filtration column (Cytivia) in size exclusion buffer: 10mM Tris, pH 7.4, 150 mM NaCl, 2mM b-mercaptoethanol and 5% glycerol. The purified VP39 was concentrated to a set of different concentrations from 7 to 45 mg/ml, mixed with sinefungin (1 mM) and directly used for crystallization trials.

### Crystallization and data collection

VP39 protein supplemented with 1 mM sinefungin and it was plated using the sitting drops method. 200 nl protein solution and 200 nl reservoir solution were mixed in several commercial crystallization screens using a mosquito robot (SPT Labtech). Only low-quality crystals were generated in the initial experiments, which were then used as crystallization seeds in further experiments. The initial crystals were crushed with a glass rod and diluted 10 000x to produce seeds. The seeding screen was prepared using Dragonfly (SPT Labtech) by gradually decreasing the PEG concentration of the original hit (200 mM lithium citrate, 20% (w/v) PEG 3350). The use of Angstrom Additive Screen TM (Molecular Dimensions) improved the quality of the seed. Finally, thin plate shaped crystals grew within 24 hours at 18°C in 200 mM lithium citrate and 14.5% (w/v) PEG 3350 and were cryoprotected in a 20% (v/v) glycerol-enriched well solution and snap-frozen in liquid nitrogen.

The crystallographic dataset was collected from a single crystal on the BL14.1 beamline at the BESSY II electron storage ring operated by the Helmholtz-Zentrum Berlin^33^. The dataset was collected at the temperature of 100 K using the wavelength of 0.9184 Å. The crystals diffracted to 2Å resolution and belonged to the P2_1_2_1_2_1_ space group. The data was integrated and scaled using XDS^34^. The structure of the MPXV VP39/sinefungin complex was solved by molecular replacement using the structure of the vaccinia virus VP39/SAH complex as a search model (pdb entry 1VP3)^35^. The initial model was obtained with Phaser^36^ from the Phenix package^37^. The model was further improved using automatic model refinement with Phenix.refine^38^ from the Phenix package^37^ and manual model building with Coot^39^. Statistics for data collection and processing, structure solution and refinement are summarized in Table 1. Final Ramachandran statistics were as follows: favored 99.64%, allowed 0.36%, outliers 0.00%. Structural figures were generated with the PyMOL Molecular Graphics System v2.0 (Schrödinger, LLC). The atomic coordinates and structural factors were deposited in the Protein Data Bank (https://www.rcsb.org) under the PDB accession code 8B07.

### RNA substrate preparation

For the 2’-O-MTase, an m7GpppG-capped RNA or an m7GpppA-capped RNA were used. These RNAs were 35 nucleotides long and they were prepared by *in vitro* transcription using a DNA template (5’-CAGTAATACGACTCACTATAGGGGAAGCGGGCATGCGGCCAGCCATAGCCGATCA-3’) and the TranscriptAid T7 high-yield transcription kit (Thermo Scientific). The reaction was performed in a 50-μl mixture containing 1× TranscriptAid reaction buffer, 7.5 mM nucleoside triphosphates [NTPs], 1 μg template DNA, and 1× TranscriptAid enzyme mix and 6 mM cap analog m7GpppG (Jena Bioscience) or m7GpppA (prepared chemically according to a published protocol^40^). The reaction was incubated at 37°C for 8 h. Next, the RNA was purified using RNA Clean & Concentrator-5 from Zymo Research. DNA was removed by DNase I treatment performed directly on the column.

### 2’-O-MTase assay

The reaction was performed in a total volume of 15 μl and all the compounds were diluted in DEPC water to prevent contamination with RNases. The reaction mixture contained 4 μM SAM (S-Adenosyl-L-methionine), 4 μM m7GpppG-capped RNA or m7GpppA-capped RNA in the MTase reaction buffer (5mM Tris pH 8.0, 1mM TCEP, 0.1 mg/ml BSA, 0.005% Triton X-100, 1mM MgCl_2_). The reaction was initiated by adding 500 nM VP39, mixed, spun, and incubated at 30°C for 120 min. The reactions were stopped by adding 30 μl of 7.5% formic acid and analyzed on an Echo mass spectrometry system coupled with a Sciex 6500 triple-quadrupole mass spectrometer operating with an electrospray ionization source. The rate of MTase activity was measured as the amount of the product of the reaction S-adenosylhomocysteine (SAH). The spectrometer was run in the multiple-reaction-monitoring (MRM) mode with the interface heated to 350°C. The declustering potential was 20 V, the entrance potential 10 V, and the collision energy 28 eV. Ten nanoliters was injected in the mobile phase (flow rate of 0.46ml/min; 70% acetonitrile with 0.1% formic acid). The characteristic product ion of SAH, m/z 385.1 >134.1, was used for quantification.

For measurements of inhibition of the 2’-O-MTase reaction by sinefungin, the above described protocol was used with only m7GpppG-capped RNA as a substrate. Sinefungin was added to the reaction mixture in the concentration range from 0 to 100 μM and all measurements were performed in triplicates, i.e. the measurements were taken from distinct sample reactions that were prepared using the same stocks of the enzyme, inhibitor, and other chemicals. The half maximal inhibitory concentration (IC_50_) was calculated by non-linear regression and curve fitting using the equation: y(x) = max/(1+x/IC 50), where y is the substrate (SAM) conversion rate, x is the concentration of inhibitor (sinefungin), and max is the maximal substrate (SAM) conversion rate in the absence of inhibitor.

## Acknowledgment

We thank the Helmholtz-Zentrum Berlin für Materialien und Energie for the allocation of synchrotron radiation beamtime. This research was funded by the project the National Institute Virology and Bacteriology (Programme EXCELES, Project No. LX22NPO5103) - Funded by the European Union - Next Generation EU. The Academy of Sciences of the Czech Republic (RVO: 61388963) is also acknowledged.

## References

1 Rizk, J. G., Lippi, G., Henry, B. M., Forthal, D. N. & Rizk, Y. Prevention and Treatment of Monkeypox. Drugs 82, 957–963, doi:10.1007/s40265-022-01742-y (2022).

2 Beer, E. M. & Rao, V. B. A systematic review of the epidemiology of human monkeypox outbreaks and implications for outbreak strategy. PLoS neglected tropical diseases 13, e0007791, doi:10.1371/journal.pntd.0007791 (2019).

3 Prevention, C. f. D. C. a. Monkeypox, https://www.cdc.gov/poxvirus/monkeypox/index.html> (2022).

4 Grimm, C., Bartuli, J. & Fischer, U. Cytoplasmic gene expression: lessons from poxviruses. Trends in biochemical sciences, doi:10.1016/j.tibs.2022.04.010 (2022).

5 Paterson, B. M. & Rosenberg, M. Efficient translation of prokaryotic mRNAs in a eukaryotic cell-free system requires addition of a cap structure. Nature 279, 692–696, doi:10.1038/279692a0 (1979).

6 Both, G. W., Furuichi, Y., Muthukrishnan, S. & Shatkin, A. J. Ribosome binding to reovirus mRNA in protein synthesis requires 5’ terminal 7-methylguanosine. Cell 6, 185–195, doi:10.1016/0092-8674(75)90009-4 (1975).

7 Mears, H. V. & Sweeney, T. R. Better together: the role of IFIT protein-protein interactions in the antiviral response. J Gen Virol 99, 1463–1477, doi:10.1099/jgv.0.001149 (2018).

8 Thoresen, D., Wang, W., Galls, D., Guo, R., Xu, L. & Pyle, A. M. The molecular mechanism of RIG-I activation and signaling. Immunol Rev 304, 154–168, doi:10.1111/imr.13022 (2021).

9 Liu, S. W., Katsafanas, G. C., Liu, R., Wyatt, L. S. & Moss, B. Poxvirus decapping enzymes enhance virulence by preventing the accumulation of dsRNA and the induction of innate antiviral responses. Cell host & microbe 17, 320–331, doi:10.1016/j.chom.2015.02.002 (2015).

10 Georgana, I., Sumner, R. P., Towers, G. J. & Maluquer de Motes, C. Virulent Poxviruses Inhibit DNA Sensing by Preventing STING Activation. Journal of virology 92, doi:10.1128/JVI.02145-17 (2018).

11 Shuman, S. RNA capping: progress and prospects. Rna 21, 735–737, doi:10.1261/rna.049973.115 (2015).

12 Hyde, J. L. & Diamond, M. S. Innate immune restriction and antagonism of viral RNA lacking 2-O methylation. Virology 479-480, 66–74, doi:10.1016/j.virol.2015.01.019 (2015).

13 Hodel, A. E., Gershon, P. D., Shi, X. N. & Quiocho, F. A. The 1.85 angstrom structure of vaccinia protein VP39: A bifunctional enzyme that participates in the modification of both mRNA ends. Cell 85, 247–256, doi:Doi 10.1016/S0092-8674(00)81101-0 (1996).

14 Krafcikova, P., Silhan, J., Nencka, R. & Boura, E. Structural analysis of the SARS-CoV-2 methyltransferase complex involved in RNA cap creation bound to sinefungin. Nature communications 11, 3717, doi:10.1038/s41467-020-17495-9 (2020).

15 Benoni, R., Krafcikova, P., Baranowski, M. R., Kowalska, J., Boura, E. & Cahova, H. Substrate Specificity of SARS-CoV-2 Nsp10-Nsp16 Methyltransferase. Viruses 13, 1722, doi:10.3390/v13091722 (2021).

16 Hodel, A. E., Gershon, P. D. & Quiocho, F. A. Structural basis for sequence-nonspecific recognition of 5’-capped mRNA by a cap-modifying enzyme. Molecular cell 1, 443–447, doi:10.1016/s1097-2765(00)80044-1 (1998).

17 Li, Z. & Lazaridis, T. Water at biomolecular binding interfaces. Physical Chemistry Chemical Physics 9, 573–581, doi:10.1039/b612449f (2007).

18 Samways, M. L., Taylor, R. D., Macdonald, H. E. B. & Essex, J. W. Water molecules at protein-drug interfaces: computational prediction and analysis methods. Chemical Society Reviews 50, 9104–9120, doi:10.1039/d0cs00151a (2021).

19 Kmiec, D. & Kirchhoff, F. Monkeypox: A New Threat? Int J Mol Sci 23, doi:10.3390/ijms23147866 (2022).

20 Bunge, E. M., Hoet, B., Chen, L., Lienert, F., Weidenthaler, H., Baer, L. R. & Steffen, R. The changing epidemiology of human monkeypox-A potential threat? A systematic review. PLoS neglected tropical diseases 16, e0010141, doi:10.1371/journal.pntd.0010141 (2022).

21 Prevention, C. f. D. C. a. JYNNEOS Vaccine. (2022).

22 Russo, A. T., Grosenbach, D. W., Chinsangaram, J., Honeychurch, K. M., Long, P. G., Lovejoy, C., Maiti, B., Meara, I. & Hruby, D. E. An overview of tecovirimat for smallpox treatment and expanded anti-orthopoxvirus applications. Expert Rev Anti Infect Ther 19, 331–344, doi:10.1080/14787210.2020.1819791 (2021).

23 Decroly, E., Ferron, F., Lescar, J. & Canard, B. Conventional and unconventional mechanisms for capping viral mRNA. Nat Rev Microbiol 10, 51–65, doi:10.1038/nrmicro2675 (2011).

24 Brecher, M., Chen, H., Li, Z., Banavali, N. K., Jones, S. A., Zhang, J., Kramer, L. D. & Li, H. Identification and Characterization of Novel Broad-Spectrum Inhibitors of the Flavivirus Methyltransferase. ACS Infect Dis 1, 340–349, doi:10.1021/acsinfecdis.5b00070 (2015).

25 Lim, S. P., Sonntag, L. S., Noble, C., Nilar, S. H., Ng, R. H., Zou, G., Monaghan, P., Chung, K. Y., Dong, H. P., Liu, B. P., Bodenreider, C., Lee, G., Ding, M., Chan, W. L., Wang, G., Jian, Y. L., Chao, A. T., Lescar, J., Yin, Z., Vedananda, T. R., Keller, T. H. & Shi, P. Y. Small Molecule Inhibitors That Selectively Block Dengue Virus Methyltransferase. Journal of Biological Chemistry 286, 6233–6240, doi:10.1074/jbc.M110.179184 (2011).

26 Devkota, K., Schapira, M., Perveen, S., Khalili Yazdi, A., Li, F., Chau, I., Ghiabi, P., Hajian, T., Loppnau, P., Bolotokova, A., Satchell, K. J. F., Wang, K., Li, D., Liu, J., Smil, D., Luo, M., Jin, J., Fish, P. V., Brown, P. J. & Vedadi, M. Probing the SAM Binding Site of SARS-CoV-2 Nsp14 In Vitro Using SAM Competitive Inhibitors Guides Developing Selective Bisubstrate Inhibitors. SLAS Discov, 24725552211026261, doi:10.1177/24725552211026261 (2021).

27 Khalili Yazdi, A., Li, F., Devkota, K., Perveen, S., Ghiabi, P., Hajian, T., Bolotokova, A. & Vedadi, M. A High-Throughput Radioactivity-Based Assay for Screening SARS-CoV-2 nsp10-nsp16 Complex. SLAS Discov 26, 757–765, doi:10.1177/24725552211008863 (2021).

28 Otava, T., Sala, M., Li, F., Fanfrlik, J., Devkota, K., Perveen, S., Chau, I., Pakarian, P., Hobza, P., Vedadi, M., Boura, E. & Nencka, R. The Structure-Based Design of SARS-CoV-2 nsp14 Methyltransferase Ligands Yields Nanomolar Inhibitors. ACS Infect Dis 7, 2214–2220, doi:10.1021/acsinfecdis.1c00131 (2021).

29 Klima, M., Khalili Yazdi, A., Li, F., Chau, I., Hajian, T., Bolotokova, A., Kaniskan, H. U., Han, Y., Wang, K., Li, D., Luo, M., Jin, J., Boura, E. & Vedadi, M. Crystal structure of SARS-CoV-2 nsp10-nsp16 in complex with small molecule inhibitors, SS148 and WZ16. Protein science: a publication of the Protein Society 31, e4395, doi:10.1002/pro.4395 (2022).

30 Ahmed-Belkacem, R., Hausdorff, M., Delpal, A., Sutto-Ortiz, P., Colmant, A. M. G., Touret, F., Ogando, N. S., Snijder, E. J., Canard, B., Coutard, B., Vasseur, J. J., Decroly, E. & Debart, F. Potent Inhibition of SARS-CoV-2 nsp14 N7-Methyltransferase by Sulfonamide-Based Bisubstrate Analogues. Journal of medicinal chemistry 65, 6231–6249, doi:10.1021/acs.jmedchem.2c00120 (2022).

31 Nencka, R., Silhan, J., Klima, M., Otava, T., Kocek, H., Krafcikova, P. & Boura, E. Coronaviral RNA-methyltransferases: function, structure and inhibition. Nucleic Acids Res 50, 635–650, doi:10.1093/nar/gkab1279 (2022).

32 Hercik, K., Brynda, J., Nencka, R. & Boura, E. Structural basis of Zika virus methyltransferase inhibition by sinefungin. Archives of virology 162, 2091–2096, doi:10.1007/s00705-017-3345-x (2017).

33 Mueller, U., Forster, R., Hellmig, M., Huschmann, F. U., Kastner, A., Malecki, P., Puhringer, S., Rower, M., Sparta, K., Steffien, M., Uhlein, M., Wilk, P. & Weiss, M. S. The macromolecular crystallography beamlines at BESSY II of the Helmholtz-Zentrum Berlin: Current status and perspectives. Eur Phys J Plus 130, doi:ARTN 14110.1140/epjp/i2015-15141-2 (2015).

34 Kabsch, W. Xds. Acta crystallographica. Section D, Biological crystallography 66, 125–132, doi:10.1107/S0907444909047337 (2010).

35 Hodel, A. E., Gershon, P. D., Shi, X. N., Wang, S. M. & Quiocho, F. A. Specific protein recognition of an mRNA cap through its alkylated base. Nature structural biology 4, 350–354, doi:DOI 10.1038/nsb0597-350 (1997).

36 McCoy, A. J., Grosse-Kunstleve, R. W., Adams, P. D., Winn, M. D., Storoni, L. C. & Read, R. J. Phaser crystallographic software. Journal of applied crystallography 40, 658–674, doi:10.1107/S0021889807021206 (2007).

37 Liebschner, D., Afonine, P. V., Baker, M. L., Bunkoczi, G., Chen, V. B., Croll, T. I., Hintze, B., Hung, L. W., Jain, S., McCoy, A. J., Moriarty, N. W., Oeffner, R. D., Poon, B. K., Prisant, M. G., Read, R. J., Richardson, J. S., Richardson, D. C., Sammito, M. D., Sobolev, O. V., Stockwell, D. H., Terwilliger, T. C., Urzhumtsev, A. G., Videau, L. L., Williams, C. J. & Adams, P. D. Macromolecular structure determination using X-rays, neutrons and electrons: recent developments in Phenix. Acta Crystallogr D 75, 861–877, doi:10.1107/S2059798319011471 (2019).

38 Afonine, P. V., Poon, B. K., Read, R. J., Sobolev, O. V., Terwilliger, T. C., Urzhumtsev, A. & Adams, P. D. Real-space refinement in PHENIX for cryo-EM and crystallography. Acta Crystallogr D 74, 531–544, doi:10.1107/S2059798318006551 (2018).

39 Emsley, P., Lohkamp, B., Scott, W. G. & Cowtan, K. Features and development of Coot. Acta crystallographica. Section D, Biological crystallography 66, 486–501, doi:10.1107/S0907444910007493 (2010).

40 Baranowski, M. R., Nowicka, A., Rydzik, A. M., Warminski, M., Kasprzyk, R., Wojtczak, B. A., Wojcik, J., Claridge, T. D., Kowalska, J. & Jemielity, J. Synthesis of fluorophosphate nucleotide analogues and their characterization as tools for (1)(9)F NMR studies. The Journal of organic chemistry 80, 3982–3997, doi:10.1021/acs.joc.5b00337 (2015).

